# Age differences in functional connectivity and dedifferentiation of category representations

**DOI:** 10.1101/2024.01.04.574135

**Authors:** Claire Pauley, Dagmar Zeithamova, Myriam C. Sander

## Abstract

With advancing age, the distinctiveness of neural representations of information declines. While the finding of this so-called ‘age-related neural dedifferentiation’ in category-selective neural regions is well-described, how neural dedifferentiation manifests at the level of large-scale functional networks is less understood. Furthermore, the relationship between age-related changes in network organization and dedifferentiation is unknown. Here, we investigated age-related neural dedifferentiation of category-selective regions as well as whole-brain functional networks. We additionally examined age differences in connectivity of category-selective regions to the rest of the brain. Younger and older adults viewed blocks of face and house stimuli while performing memory encoding and retrieval in the fMRI scanner. We found an age-related decline in neural distinctiveness for faces in the fusiform gyrus (FG) and for houses in the parahippocampal gyrus (PHG). Functional connectivity analyses revealed age-related dedifferentiation of global network structure as well as age differences in the connectivity profiles to category-selective regions. Together, our findings suggest that age-related neural dedifferentiation manifests both in regional categorical representations as well as in whole-brain functional networks.

**Highlights:**

- Category representations are less distinctive, or dedifferentiated, in older adults
- Functional networks are less segregated in older adults
- Older adults reveal less connectivity between fusiform gyrus and visual cortices

## 1. Introduction

Aging often results in declines in various cognitive functions, including episodic memory. Age differences in episodic memory performance have been related to the way information is neurally represented, suggesting that neural responses are less distinctive in older adults as compared to younger adults – a phenomenon called “age-related dedifferentiation” (for reviews, see Koen et al., 2020; Koen & Rugg, 2019; Sommer & Sander, 2022). At the level of processing within individual regions-of-interest (ROIs), this reduction in distinctiveness is often investigated in terms of an age-related decline in the selectivity of high-level visual cortices for particular stimulus categories.

Senescent declines in neural distinctiveness have been reported in the parahippocampal gyrus (PHG) during scene processing (Koen, 2022; Koen et al., 2019; Srokova et al., 2020), in the fusiform gyrus (FG) for faces (Park et al., 2004, 2012; Pauley et al., 2023), in lateral occipital cortex for objects (Chee et al., 2006; Koen, 2022), and in the visual word form area for words (Park et al., 2004; but, see Koen et al., 2019; Payer et al., 2006 for null age effects).

The term “age-related dedifferentiation” has also been utilized in the context of age differences observed in functional network structures (Koen et al., 2020). Studies assessing age differences in functional connectivity frequently report less segregated network structures in older adults, characterized by a reduction in within-network connectivity as well as an increase in between-network connectivity both in pre-defined networks and in networks derived through a graph-theoretical framework (Betzel et al., 2014; Cao et al., 2014; Chan et al., 2017, 2014; Geerligs et al., 2015; Iordan et al., 2017; King et al., 2018). These age-related reductions in segregation across large-scale functional brain networks are also referred to as network-level dedifferentiation (Koen et al., 2020). One of the most consistent findings across studies is the negative impact of age on the segregation of the default mode network with several findings of reduced within-default-mode network communication in older adults (Andrews-Hanna et al., 2007; Betzel et al., 2014; Geerligs et al., 2015; Mak et al., 2017; Song et al., 2014). Senescent declines in default mode connectivity have previously been associated with declines in longitudinal recognition memory (Persson et al., 2014), indicating a link between age-related network reorganization and memory performance. The link between memory and connectivity was further supported by Chan and colleagues (2014), who identified a similar cross-sectional relationship between memory performance and segregation of association (non-sensory) functional networks. In sum, aging is associated with a decline in the specificity of functional network architecture, driven by both attenuated within-network connectivity as well as increased between-network connectivity, that may be tied to senescent memory decline.

So far, little is known about the mechanisms behind the observed interindividual variability in neural distinctiveness. Interestingly, in younger adults, one line of research examined the connectivity of category-selective ROIs as a potential mechanism driving categorical distinctiveness. Connectivity patterns to category-selective ROIs have been used to decode the preferred stimulus category (Chen et al., 2017; Wang et al., 2016) and have been shown to flexibly adapt to the attended stimulus category (Córdova et al., 2016; Keller et al., 2022; Norman-Haignere et al., 2012; Silson et al., 2019). For example, Norman-Haignere and colleagues (2012) found that connectivity between the early visual cortex and ventral temporal cortices varied depending on whether participants were directing their attention to scenes or faces. Specifically, when individuals paid attention to scenes, the connectivity between the PHG and calcarine cortex increased, whereas when they were attending faces, the connectivity between the FG and calcarine cortex increased. Thus, connectivity patterns to category-selective ROIs may support category representations within these regions. Despite these findings, it is still unknown whether there are age differences in connectivity to category-selective ROIs.

In this study, we were first interested in the influence of age on neural distinctiveness on categorical representations (i.e., regional dedifferentiation) as well as on large-scale functional networks (i.e., network dedifferentiation). Crucially, we additionally investigated how age impacted connectivity patterns to category-selective regions. Finally, we asked how network segregation and connectivity to category-selective regions were related to distinctiveness of categorical representations. Using a memory paradigm in which younger and older adults were presented with blocks of face and house stimuli, we measured age differences in neural distinctiveness of face and house processing in category-selective visual cortices (FG and PHG) with multi-voxel pattern analysis. We further assessed age differences in global network segregation as well as in connectivity between category-selective regions and global functional networks. Using correlation analyses, we investigated the link between regional and network dedifferentiation as well as the relationship between regional dedifferentiation and the connectivity patterns to these regions.

## 2. Materials and Methods

Data from this project were previously reported in Pauley et al. (2023). Methods and analyses that are relevant for the current study are repeated here. Two additional papers based on this dataset (Kobelt et al., 2021; Pauley et al., 2022) were later retracted by the authors due to a preprocessing error. The retracted papers (and corrected findings) can be found at https://osf.io/t8dpv/ and https://osf.io/7n3mz/.

### 2.1 Participants

Data were collected from a total of 76 healthy adults. Participants were recruited within 2 age groups: younger adults (18–27 years, N = 39) and older adults (64–76 years, N = 37). Four participants were excluded due to excessive motion in the scanner (1 younger adult and 3 older adults; see Section 2.5 for details), 3 were excluded due to memory performance below chance level (2 younger adults and 1 older adult), and 2 were excluded due to poor MRI data quality (1 younger adult and 1 older adult).

The final sample consisted of 35 younger adults (M(SD) age = 22.3 (2.7) years, 16 females, 19 males) and 32 older adults (M(SD) age = 70.8 (2.5) years, 18 females, 14 males). Participants were screened via telephone for mental and physical illness, metal implants, and current medications. Additionally, all older adults were screened using the Mini-Mental State Examination (Folstein et al., 1975) and all exceeded the threshold of 26 points. The study was approved by the ethics committee of the German Society for Psychological Research (DGPs) and written informed consent was obtained from each participant at the time of the study.

### 2.2 Stimuli

Stimuli were comprised of 300 grayscale images belonging to 3 different categories: 120 neutral faces (adapted from the FACES database; Ebner et al., 2010), 120 houses (some obtained online and some adapted from Park et al., 2004), and 60 phase-scrambled images (30 faces and 30 houses, constructed from randomly selected face and house images) serving as control stimuli. An additional image from each category was selected to serve as target stimuli for the encoding target-detection task (see Section 2.3). All nontarget face and house images were randomly divided into 2 sets of 120 images (60 faces and 60 houses). One stimulus set was presented during both encoding and recognition (old images) and the other set was presented only during recognition (new images). The same stimulus sets were used for all participants.

### 2.3 Experimental design

The following paradigm was part of a larger study spanning 2 days of data collection. This study focuses only on the face-house task, which comprised an incidental encoding phase and a surprise recognition test, both conducted inside the fMRI scanner on the same day with a delay of approximately 30 min (see Figure 1). The encoding phase consisted of 2 identical runs each with 9 stimulus blocks. In order to ensure the participants were paying attention to the stimuli, they were asked to perform a target-detection task in which they pressed a button when 1 of 3 pre-learned target images was presented. Stimuli were randomly distributed into the blocks such that each block contained 20 images of a single category (faces, houses, or phase-scrambled) as well as a category-matched target image. The block order was alternating and counterbalanced across participants, always starting with either a face or house block. The stimulus order within each block was pseudo-randomized with the condition that the target image was not presented in either the first 4 or last 4 trials of a block. Due to a technical problem, the same stimulus order was used for all participants who started with a face block and for 36 of the participants starting with a house block. Prior to the encoding phase, participants completed 5 practice trials of each stimulus category, including each of the target stimuli, to verify that they understood the target-detection task. The non-target training stimuli were excluded from the main experiment. Since the 2 encoding runs were identical, participants were exposed to each stimulus twice during the encoding phase. Phase-scrambled images were not used in any subsequent analyses in this project. Stimuli were presented for 1200 ms and separated by a fixation cross with a jittered duration between 500 and 8000 ms. In total, the encoding phase lasted approximately 22 min.

**Figure 1.**
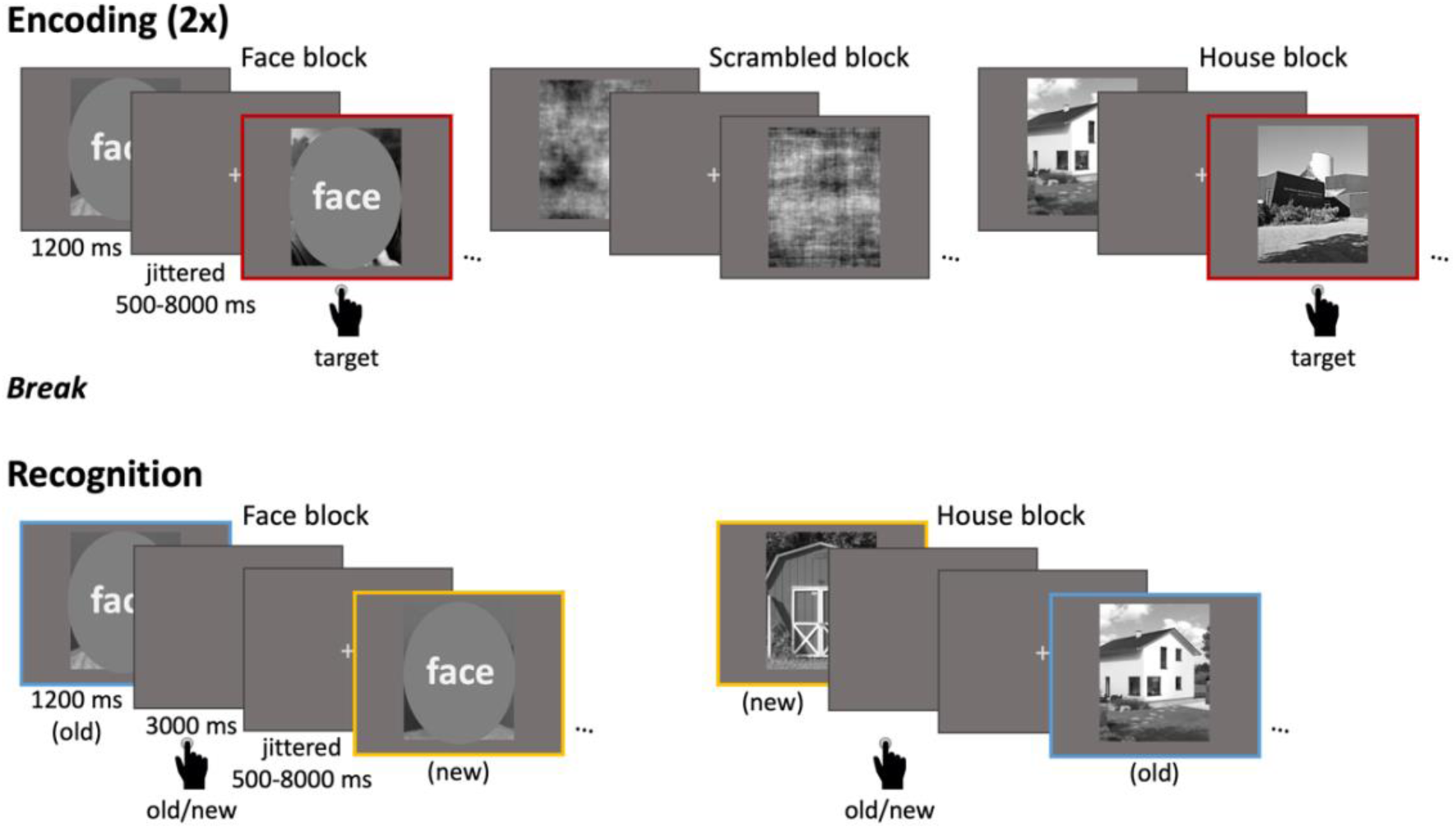
Face-house task design. This fMRI paradigm comprised an incidental encoding phase (top) and a surprise recognition test (bottom). During encoding, two identical runs of face, house, and phase-scrambled images (not assessed here) were presented in a block design with 9 stimulus blocks each (3 alternating blocks from each stimulus category). Each block had 21 trials (20 exemplars of the respective category and 1 pre-learned target stimulus). Participants were instructed to press a button when a target stimulus appeared. During the recognition test, six alternating face and house blocks were presented with 40 trials each (20 old trials from encoding and 20 new trials). Participants indicated via button press whether each image was old or new. Figure reproduced from Pauley et al. (2023).

Following encoding, participants remained in the scanner briefly while structural scans were collected (see below). Then, they had a short break outside the scanner while they received instructions for the recognition test. They then returned to the scanner to complete the recognition test. The recognition test consisted of 6 blocks (3 face and 3 house) presented across 2 functional runs (3 blocks per run), alternating between face and house blocks. Instructions explaining the task were presented for 7 s at the beginning of each scanning run. Each block contained 20 old images (seen during encoding) and 20 new images of the same stimulus category. For each trial, participants were asked whether the image was old or new, which they indicated via button press. The stimulus order was pseudorandomized such that no more than 3 old or new images were presented consecutively.

Due to a technical problem, the same stimulus order was used for 13 participants who started with a face block and 14 participants who started with a house block. Stimuli were presented for 1200 ms and followed by a gray screen for 3000 ms in which participants could give their response. Fixation crosses separated the trials with jittered durations between 500 and 8000 ms. In total, the recognition task lasted approximately 26 min.

### 2.4 Behavioral data analyses

Since additional participants were excluded here compared with Pauley et al. (2023), we replicated the behavioral analyses initially reported. Recognition memory performance was assessed as the difference between the hit rate (proportion of correctly identified old stimuli) and the false alarm rate (proportion of new stimuli incorrectly identified as old stimuli (Snodgrass and Corwin, 1988).

Participants with recognition memory performance less than zero, indicating a higher probability of responding “old” to new stimuli as opposed to old stimuli, were excluded from subsequent analyses (see also Section 2.1). Dependent-samples *t*-tests were used to determine whether memory performance exceeded chance level. A two-way mixed factorial analysis of variance (ANOVA) was used to assess age differences in recognition memory performance as well as differences related to stimulus type (face vs. house).

### 2.5 fMRI data acquisition and preprocessing

Brain imaging was acquired with a Siemens Magnetom TrioTim 3T MRI scanner with a 32-channel head-coil. Functional images were collected using an echo planar imaging sequence during both the encoding and recognition phases in 2 runs each. Each encoding run consisted of 270 volumes and each recognition run consisted of 372 volumes (voxel size = 3 × 3 × 3 mm^3^; slice gap = 0.3 mm; TR = 2 s; TE = 30 ms). The first 3 volumes of each run were dummy volumes and were excluded prior to preprocessing. Following the encoding phase, a T1-weighted (T1w) magnetization prepared rapid acquisition gradient echo (MPRAGE) pulse sequence image was acquired (voxel size = 1 × 1 × 1 mm^3^; TR = 2.5 ms; TE = 4.77 ms; flip angle = 7°; TI = 1.1 ms). Additionally, turbo spin-echo proton density images, diffusion tensor images, and fluid attenuation inversion recovery images were collected, but not included in the following analyses. Experimental stimuli, which participants viewed via a mirror mounted on the head-coil, were projected using the Psychtoolbox (Psychophysics Toolbox) for MATLAB (Mathworks Inc., Natick, MA).

MRI data were organized according to the Brain Imaging Data Structure (BIDS) specification (Gorgolewski et al., 2016) and preprocessed using fMRIPrep (version 1.4.0; Esteban et al., 2019) with the default settings. The T1w image was corrected for intensity nonuniformity, skull-stripped, and spatially normalized to the ICBM 152 Nonlinear Asymmetrical template version 2009c through nonlinear registration. Functional images were motion-corrected, slice-time corrected, and co-registered to the normalized T1w reference image. No spatial smoothing was applied. As mentioned in Section 2.1, 4 participants were excluded due to excessive motion in the scanner. Two rounds of excessive motion detection were performed. First, two participants were excluded due to having multiple (more than one) motion artifacts within a given scanner run resulting in framewise displacements greater than the size of the voxel (3 mm; see Power et al., 2012). Second, two additional participants were excluded for having three functional runs each with more than 15% of volumes with framewise displacements > 0.5 mm or DVARS > 0.5%.

### 2.6 ROIs

Category-selective regions were defined according to the Automated Anatomical Labeling (AAL) atlas (Version 2; Tzourio-Mazoyer et al., 2002). We isolated the FG and PHG because of their functional roles in face and house processing, respectively (Epstein and Kanwisher, 1998; Kanwisher et al., 1997). Since the anterior FG and PHG have been shown to respond to stimulus categories other than faces and houses (Barense et al., 2010), the FG and PHG were manually segmented to include only the posterior portions of these regions. This was implemented by dividing the number of slices in the *y* dimension by two and manually selecting the posterior half of these slices for each ROI. We used the Schaefer atlas (7 networks, 100 parcels; Schaefer et al., 2018) for connectivity analyses.

Schaefer parcels that overlapped with the category-selective regions were excluded.

### 2.7 Multi-voxel pattern analysis to assess age differences in neural distinctiveness in category-selective ROIs

We were particularly interested in age differences in neural distinctiveness in regions particularly selective for our visual categories (i.e., faces and houses). Therefore, we measured neural specialization for faces in the posterior FG and for houses in the posterior PHG. First, event-related generalized linear models (GLMs) were performed separately for encoding and recognition. For encoding, four event types were modeled: houses, faces, phase-scrambled images, and target images. For recognition, three event types were modeled: houses, faces, and the instructional period at the beginning of each run. All trials were modeled using finite impulse response (FIR) basis sets, beginning at trial onset and lasting for 20 s post-trial onset for encoding trials and 24 s for recognition trials (for similar methodology, see Wolosin et al., 2012). In each model, 36 motion regressors were additionally included according to Power et al. (2012): six motion parameters, cerebrospinal fluid signal, white matter signal, global signal, and each of their derivatives. The models further included high-pass filtering with a 100s window. The beta maps corresponding to 2-8 s post-stimulus onset from the event-related GLMs (described above) were averaged for each stimulus. Regional distinctiveness was measured separately for encoding and recognition and was defined as the difference between within-category similarity and between-category similarity (see Haxby et al., 2001 for similar methodology). Within-category similarity was defined as the across-voxel Pearson correlation of a particular stimulus category in the first run (or either encoding or recognition) to the same stimulus category in the second run (e.g., face-face similarity). Between-category similarity was defined as the across-voxel Pearson correlation of a particular stimulus category in the first run to the other stimulus category in the second run (i.e., face-house similarity). In the FG, within-category similarity was considered face-face similarity, whereas in the PHG, within-category similarity was considered house-house similarity. Correlation coefficients were Fisher-*z* transformed before taking the difference of within- and between-category similarity. We first checked that regional distinctiveness values in the FG and PHG were reliably above zero using dependent-samples *t*-tests. Age differences in regional distinctiveness were evaluated using a 2 (age group) x 2 (ROI: FG/PHG) x 2 (memory phase: encoding/recognition) mixed factorial ANOVA on distinctiveness values (i.e., within-category – between-category similarity). Significant interactions were further evaluated using pairwise comparisons.

### 2.8 Functional connectivity analysis

In order to utilize our task-based functional data to derive connectivity as well as distinctiveness measures, we employed “background” connectivity, in which task-related responses are modeled and the residual time series is used to examine connectivity. Background connectivity can be used analogously to resting state connectivity (Fair et al., 2007; Frank and Zeithamova, 2023) and, in this case, enabled us to avoid circularity problems, which could have arisen if the same functional data were used for both the connectivity and distinctiveness metrics. The resulting residual timeseries were first cleaned by removing (or “scrubbing”) volumes that exceeded framewise displacement > 0.5 mm or DVARS > 0.5% as well as each subsequent volume. Single volumes left in between scrubbed volumes as well as the first two volumes in each run were additionally removed. The timeseries cleaning resulted in removal of an average of 3.04% of volumes for younger adults and 6.71% of volumes for older adults. The scrubbed timeseries were averaged across all voxels within each ROI and correlated across ROIs using Pearson correlation coefficient separately for encoding and recognition. Correlation coefficients were than standardized using Fisher’s *r*-to-*z* transform.

### 2.9 Assessing age differences in global network segregation

Next, we were interested in whether age-related neural dedifferentiation disrupted whole-brain functional network organization. We measured age differences in whole-brain network segregation by taking the difference in mean connectivity of all ROIs within each network (i.e., within-network connectivity) and mean connectivity between all networks (i.e., between-network connectivity) as a proportion of mean within-network connectivity (see Chan et al., 2014 for similar methodology). We conducted a 2 (age group) x 2 (memory phase: encoding/recognition) ANOVA on these mean segregation scores. Significant interactions were further evaluated using independent-samples *t*-tests. To illustrate the specific ROI-ROI connections underlying age differences in network-level segregation, we additionally used a series of independent-samples *t*-tests to test for age differences in the connectivity between all ROIs. Significance was determined by FDR-corrected *p* values.

### 2.10 Assessing connectivity to category-selective ROIs and age differences therein

We first sought to understand to which networks category-selective ROIs were most connected. To this end, we averaged the connectivity scores between each category-selective ROI and each functional network (e.g., mean connectivity of FG to the visual network, averaged across all visual network ROIs). This averaging was performed independently for each memory phase and each age group. In order to determine the preferential connectivity of each category-selective ROI, we used a series of one-sample *t*-tests to determine whether the connectivity between each of the category-selective ROIs and functional networks differed from the mean connectivity of each category-selective ROI to all networks. Significance was determined by FDR-corrected *p* values. Significant preferential connectivity to category-selective ROIs was followed up by using 2 (age group) x 2 (memory phase: encoding/recognition) mixed factorial ANOVA on mean connectivity scores.

Significant interactions were followed up using mean comparisons. Due to the high congruence of connectivity (and age differences therein) between encoding and recognition, we averaged all measures across memory phase for all subsequent analyses.

### 2.11 Relating neural distinctiveness to connectivity metrics

As an exploratory analysis, we were interested in whether network segregation or connectivity to the category-selective ROIs was associated with the magnitude of regional distinctiveness across individuals.

In order to assess whether neural dedifferentiation manifests similarly across category-selective regions and functional networks, we performed zero-order Pearson correlations across all participants (as well as within each age group) to evaluate the relationship between mean distinctiveness (averaged across category-selective ROIs) and mean network segregation.

Next, we tested whether age differences in the magnitude of neural distinctiveness were associated with age differences in preferential connectivity to category-selective ROIs. To this end, we performed zero-order Pearson correlations relating the magnitude of neural distinctiveness (as defined by multi-voxel pattern analysis) to the preferential connectivity of the corresponding category-selective ROI. Specifically, since we identified age differences only in preferential connectivity between the FG and visual network, we correlated neural distinctiveness in the FG to mean FG-visual network connectivity.

In order to verify that age differences in in-scanner motion did not influence any brain-brain relationships, we controlled any significant correlations for mean framewise displacement averaged across all runs using first-order Pearson correlations.

### 2.12 Relating distinctiveness and connectivity to memory performance

Finally, we assessed how interindividual differences in neural distinctiveness and connectivity relate to memory performance. To this end, we implemented zero-order Pearson correlations within and across age groups relating memory performance to (1) neural distinctiveness (averaged across category-selective ROIs), (2) mean network segregation, and (3) mean connectivity between the FG and the visual network.

### 2.13 Data and code availability

Data and code for performing and reproducing the analyses are available on the Open Science Framework (OSF) at https://osf.io/qh8un/.

## 3. Results

### 3.1 Behavioral results

Recognition performance (hit rate – false alarm rate) exceeded chance level in both younger (*t*(34) = 11.88, *p* < 0.001, *d* = 1.96) and older adults (*t*(31) = 9.61, *p* < 0.001, *d* = 1.66). We used a mixed factorial ANOVA to test for age and stimulus differences in memory outcomes. This analysis revealed no main effect of age (*F*(1,65) = 1.79, *p* = 0.19, partial-*η*^2^ = 0.56), indicating that memory performance did not differ between younger (*M* ± *SD* = 0.24 ± 0.12) and older adults (*M* ± *SD* = 0.20 ± 0.12). Furthermore, neither main effect of stimulus type (*F*(1,65) = 3.21, *p* = 0.08, partial-*η*^2^ = 0.41) nor interaction (*F*(1,65) = 0.21, *p* = 0.65, partial-*η*^2^ = 0.03) was identified. These findings corroborate our prior analyses of this data (Pauley et al., 2023), which included three more participants.

### 3.2 Age differences in regional distinctiveness

Using *t*-tests, we first verified that regional distinctiveness values were greater than zero (all *p*s < 0.01). Then, we performed a 2 (age group) x 2 (ROI) x 2 (memory phase) ANOVA to examine age differences in regional distinctiveness values. We found a main effect of age group (*F*(1,65) = 16.79, *p* < 0.001, partial-*η*^2^ = 0.49), indicating that younger adults (*M* ± *SD* = 0.14 ± 0.15) had higher regional distinctiveness than older adults (*M* ± *SD* = 0.06 ± 0.12; see Figure 2). Furthermore, we found a main effect of ROI (*F*(1,65) = 19.01, *p* < 0.001, partial-*η*^2^ = 0.35), suggesting that distinctiveness was higher in the FG (*M* ± *SD* = 0.14 ± 0.11) than the PHG (*M* ± *SD* = 0.07 ± 0.13). No main effect of memory phase was identified (*F*(1,65) = 0.43, *p* = 0.51, partial-*η*^2^ = 0.01).

**Figure 2.**
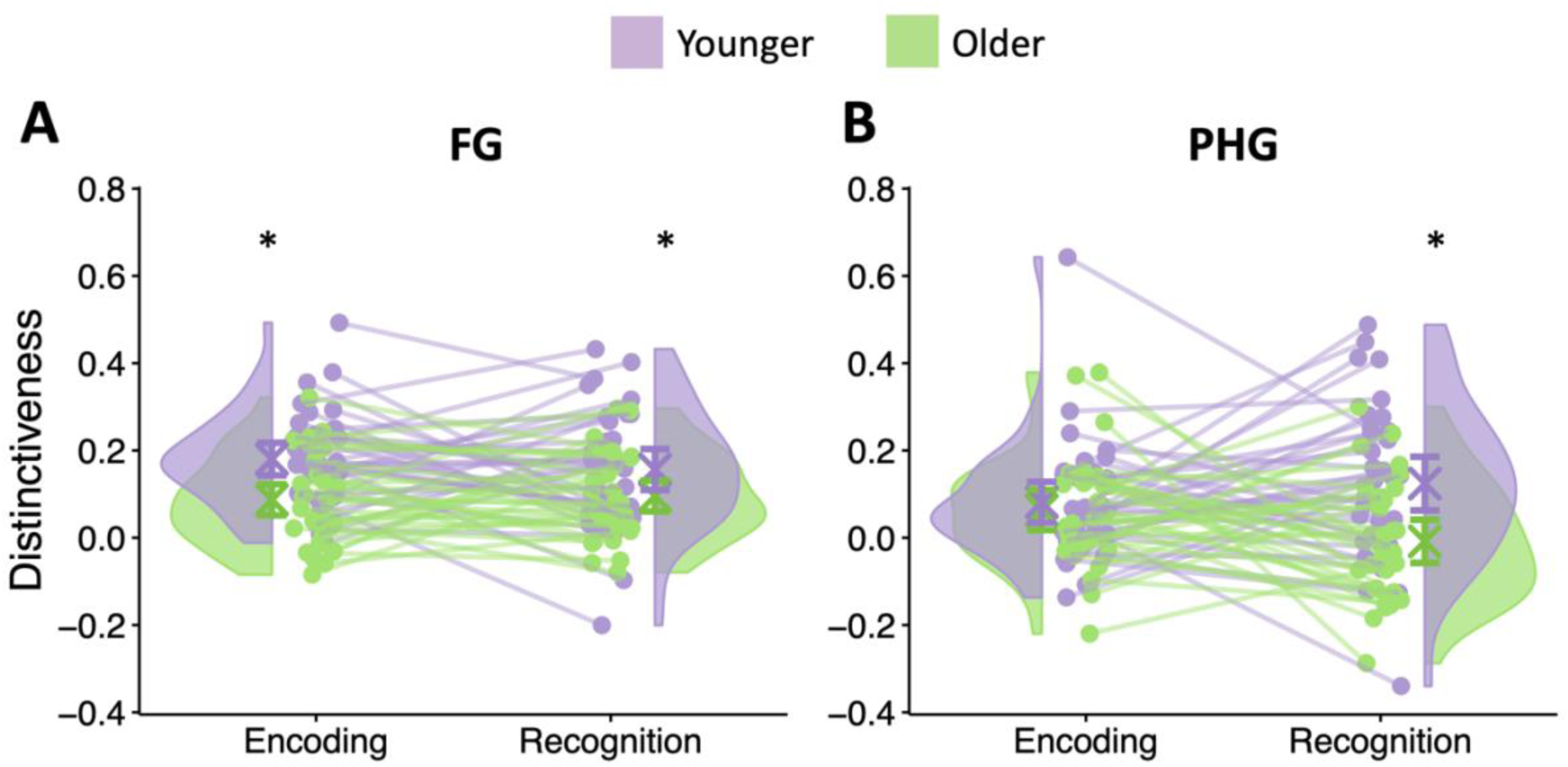
Age differences in regional distinctiveness of category-selective ROIs. Younger adults (purple) demonstrated higher regional distinctiveness (the difference between within-category and between-category similarity) than older adults (green) in the FG (A) and PHG (B) as defined by multi-voxel pattern analysis. This age effect reached significance during recognition in both the FG and PHG, however only in the FG during encoding. Half-violin plots illustrate the sample density of distinctiveness within each ROI and lines connect individual participants.

Furthermore, the interactions between age group and ROI, age group and memory phase, as well as ROI and memory phase were not significant (*p*s > 0.22). However, a three-way interaction between age group, ROI, and memory phase was identified (*F*(1,65) = 7.52, *p* = 0.008, partial-*η*^2^ = 0.11), suggesting that the age differences in distinctiveness between encoding and recognition differed between the FG and PHG. Follow-up *t*-tests revealed that younger adults exhibited greater regional distinctiveness than older adults in the FG during both encoding (*t*(65) = 3.65, *p* < 0.001, *d* = 0.88) and recognition (*t*(65) = 2.07, *p* = 0.04, *d* = 0.50) as well as in the PHG during recognition (*t*(65) = 3.37, *p* = 0.001, *d* = 0.82). No age differences in regional distinctiveness in the PHG during encoding were found (*t*(65) = 0.55, *p* = 0.58, *d* = 0.13).

### 3.3 Age-related dedifferentiation of global functional networks

Prior studies have reported that older adults demonstrate less segregated global networks characterized by a decrease in within-network connectivity and an increase in between-network connectivity (Geerligs et al., 2015). To understand whether our sample demonstrates analogous age effects, we performed a 2 (age group) x 2 (memory phase: encoding/recognition) ANOVA on mean segregation scores, defined as the difference between mean within-network connectivity and mean between-network connectivity. We found a significant main effect of age group (*F*(1,65) = 125.95, *p* < 0.001, partial-*η*^2^ = 0.95; see Figures 3 and 4), indicating that segregation was greater in younger adults (*M* ± *SD* = 1.07 ± 0.07) compared to older adults (*M* ± *SD* = 0.83 ± 0.12). We additionally found a significant main effect of memory phase (*F*(1,65) = 36.92, *p* < 0.001, partial-*η*^2^ = 0.04), revealing that segregation was greater during encoding (*M* ± *SD* = 0.98 ± 0.14) than during recognition (*M* ± *SD* = 0.93 ± 0.17). Furthermore, we found an interaction between age group and memory phase (*F*(1,65) = 15.06, *p* < 0.001, partial-*η*^2^ = 0.02). Follow-up *t*-tests revealed significant age effects in both memory phases, but with slightly greater age differences in segregation during recognition (*t*(65) = 11.71, *p* < 0.001, *d* = 2.83) than during encoding (*t*(65) = 9.34, *p* < 0.001, *d* = 2.26).

**Figure 3.**
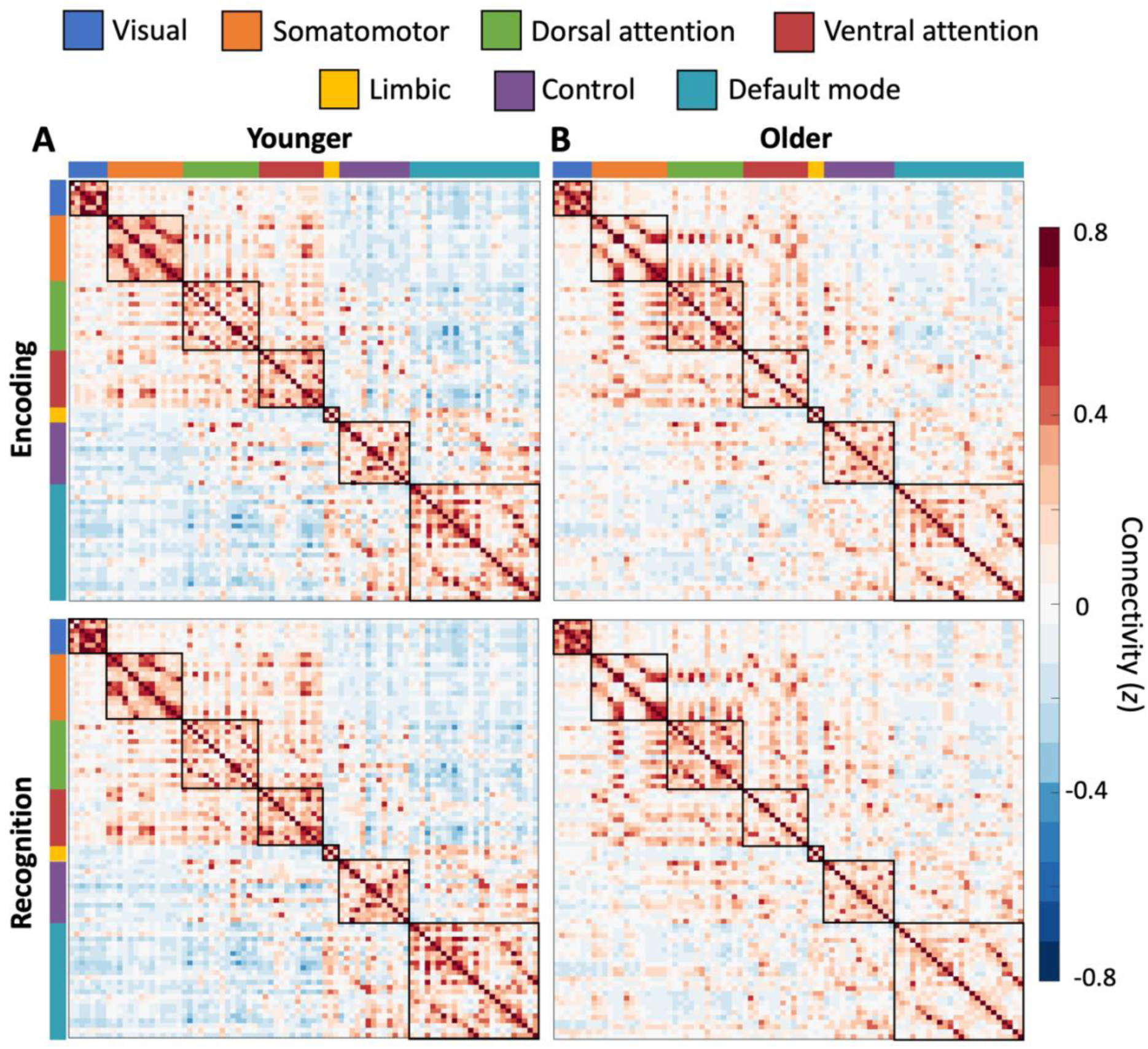
Mean ROI-to-ROI connectivity matrices for younger adults (A) and older adults (B) during encoding (top) and recognition (bottom). Black boxes outline within-network connections and networks are color-coded outside the matrices (see legend).

**Figure 4.**
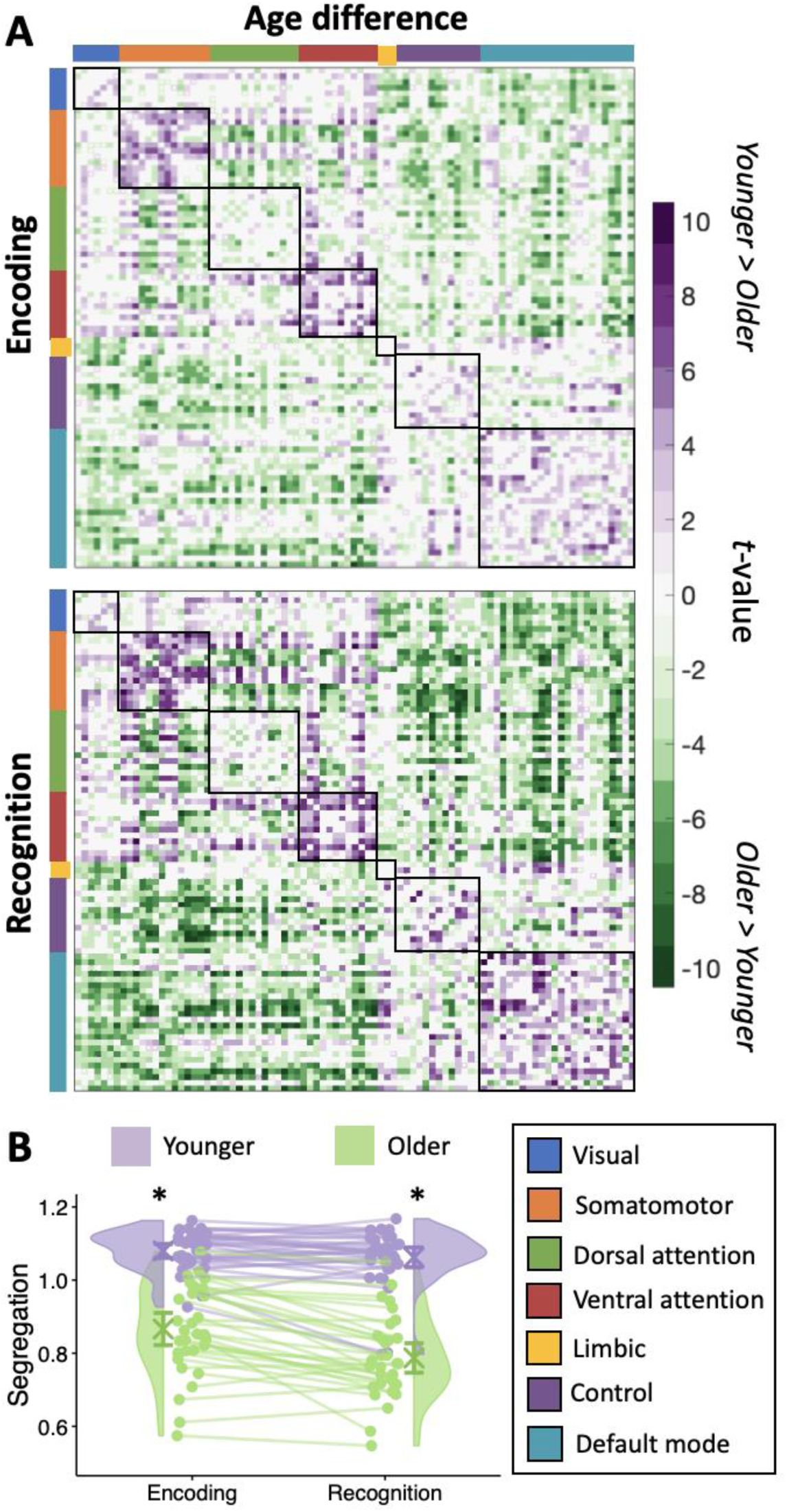
Age differences (t statistics) in ROI-to-ROI connectivity (A) during encoding (top) and during recognition (bottom). Black boxes outline within-network connections and networks are color-coded outside the matrices (see legend). Only t statistics surviving an uncorrected threshold of p < 0.05 are displayed. White spots indicate age effects that do not survive FDR-correction. Younger adults (purple) demonstrated greater overall network segregation than older adults (green; B). Half-violin plots illustrate the sample density of segregation. Error bars indicate standard error of the mean and lines connect individual participants.

### 3.4 Functional connectivity to category-selective ROIs

Here, we were interested in understanding to which networks category-selective ROIs were most connected. A series of one-sample *t*-tests revealed that the FG was preferentially connected to the visual network compared to other networks during both encoding (younger: *t*(6) = 5.53, *p* = 0.01; older: *t*(6) = 5.20, *p* = 0.01) and recognition (younger: *t*(6) = 5.56, *p* = 0.01; older: *t*(6) = 5.15, *p* = 0.01; see Figure 5). No other networks were preferentially connected to the FG (*p*s > 0.14). We were additionally interested in whether there were age differences in the preferential connectivity of the FG to the visual network. To explore this, we used a 2 (age group) x 2 (memory phase: encoding/recognition) ANOVA on mean connectivity between the FG and visual network (averaged across connectivity to all ROIs within the visual network). The ANOVA revealed a main effect of age (*F*(1,65) = 19.28, *p* < 0.001, partial-*η*^2^ = 0.80), suggesting that the FG was more strongly connected to the visual network in younger adults (*M* ± *SD* = 0.35 ± 0.10) compared to older adults (*M* ± *SD* = 0.26 ± 0.09; see Figure 6). Furthermore, a main effect of memory phase was found (*F*(1,65) = 24.05, *p* < 0.001, partial-*η*^2^ = 0.19), indicating that connectivity between the FG and the visual network was stronger during recognition (*M* ± *SD* = 0.33 ± 0.10) compared to encoding (*M* ± *SD* = 0.29 ± 0.10). No interaction between age group and memory phase (*F*(1,65) = 0.91, *p* = 0.34, partial-*η*^2^ = 0.01) was found.

**Figure 5.**
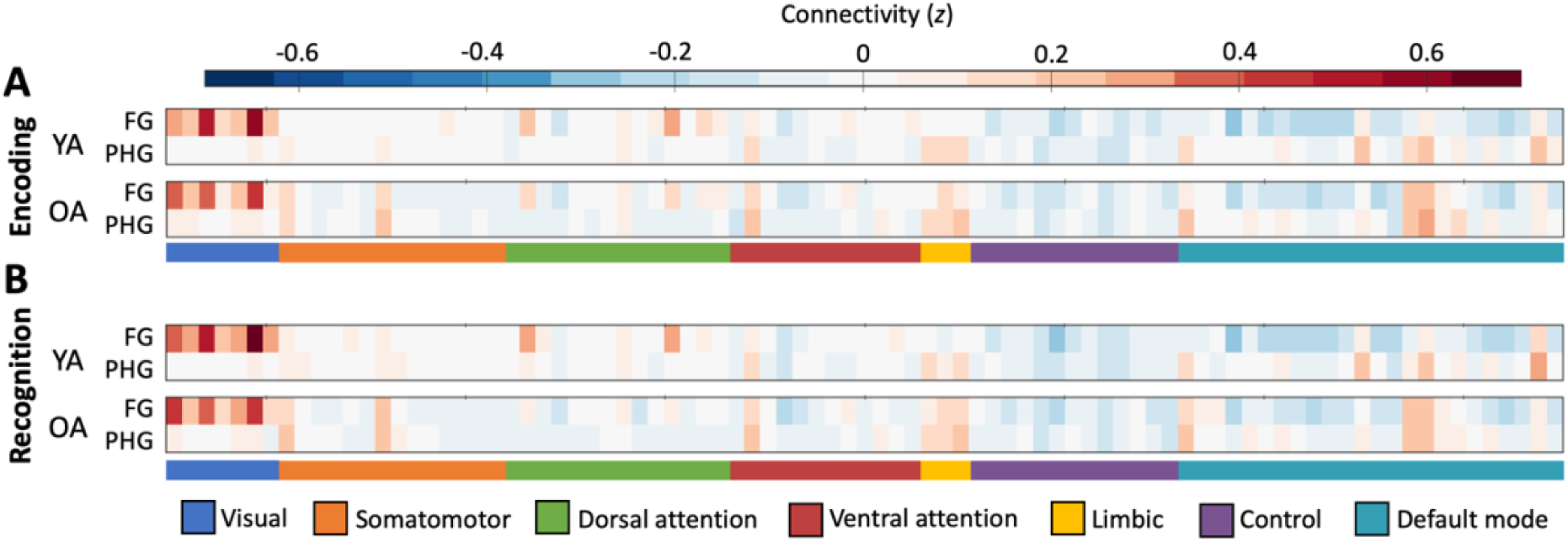
The FG was particularly connected to regions of the visual network in younger adults (YA) and older adults (OA) during both encoding (A) and recognition (B).

**Figure 6.**
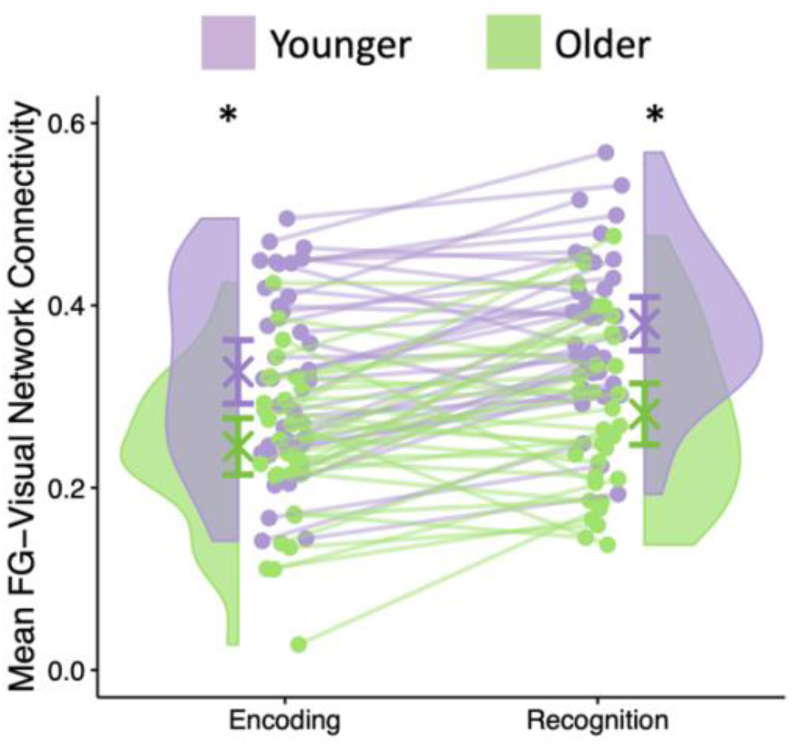
Younger adults (purple) demonstrated greater connectivity between the FG and the visual network (averaged across connectivity to all ROIs within the visual network) than older adults (green). Half-violin plots illustrate the sample density of mean connectivity. Error bars indicate standard error of the mean and lines connect individual participants.

For the PHG, a series of one-sample *t*-tests revealed that the limbic network was preferentially connected during both encoding (younger: *t*(6) = 5.14, *p* = 0.02; older: *t*(6) = 5.27, *p* = 0.01) and recognition (*t*(6) = 4.81, *p* = 0.02; older: *t*(6) = 5.26, *p* = 0.01). No other networks revealed preferential connectivity with the PHG (*p*s > 0.06), however, the control network was less connected compared to the mean in younger adults during encoding (*t*(6) = -4.00, *p* = 0.03). In order to explore age differences in the preferential connectivity of the PHG to the limbic network, we again used a 2 (age group) x 2 (memory phase: encoding/recognition) ANOVA on mean connectivity between the PHG and limbic network. The ANOVA revealed a main effect of age group (*F*(1,65) = 4.54, *p* = 0.04, partial-*η*^2^ = 0.98), ), suggesting that the PHG was more strongly connected to the limbic network in older adults (*M* ± *SD* = 0.18 ± 0.11) compared to younger adults (*M* ± *SD* = 0.14 ± 0.07). However, no main effect of memory phase (*F*(1,65) = 0.18, *p* = 0.68, partial-*η*^2^ = 0.02) as well as no interaction (*F*(1,65) = 0.01, *p* = 0.94, partial-*η*^2^ = 0.00), indicating that the preferential connectivity between the PHG and limbic network did not differ by age group or memory phase.

Due to the strong consistency in results across encoding and recognition, we averaged across memory phase for all subsequent analyses. Furthermore, since we were primarily interested in connectivity to category-selective cortices susceptible to aging, we isolated the connectivity between the FG and visual network for further analyses.

### 3.5 Age-related association between regional dedifferentiation and network dedifferentiation

We were further interested in whether the magnitude of regional distinctiveness (averaged across category-selective ROIs) was related to interindividual variation in network segregation. Pearson correlations revealed a positive relationship between distinctiveness and segregation across the whole sample (*r* = 0.42, *p* < 0.001; see Figure 7), even after controlling for framewise displacement (partial *r* = 0.30, *p* = 0.01). However, no within-group relationships reached significance (*p*s > 0.24), indicating that this relationship was driven by age differences in both distinctiveness and segregation.

**Figure 7.**
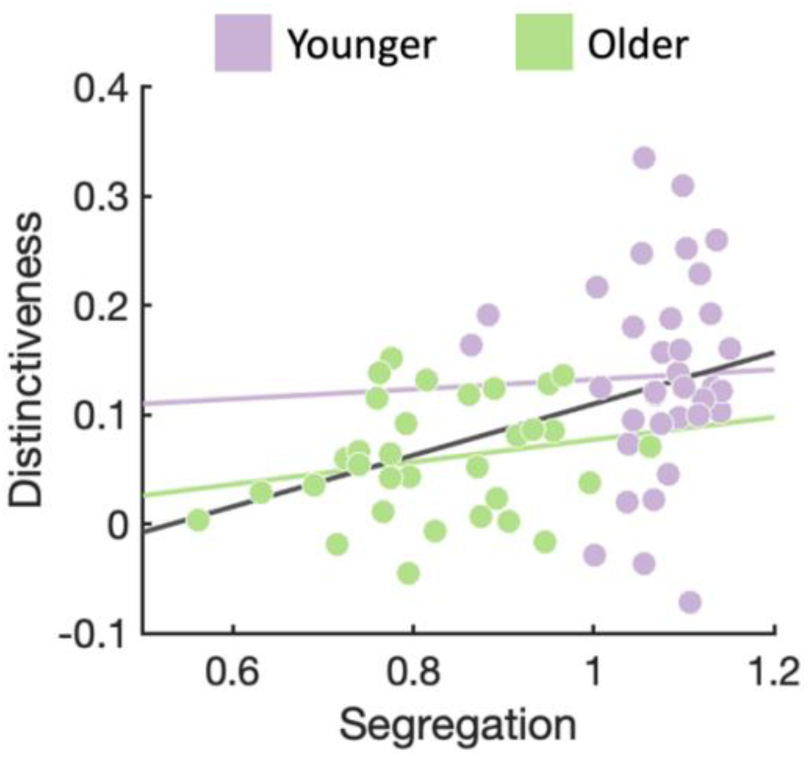
Scatter plot showing mean network segregation and mean distinctiveness (averaged across the FG and PHG). Younger adults are shown in purple, older adults in green, and the whole sample in gray.

### 3.6 No association between regional distinctiveness and connectivity to category-selective ROIs

Our analyses investigating connectivity to category-selective ROIs uncovered age differences in connectivity between the FG and visual network, with younger adults demonstrating greater connectivity than older adults, as well as age differences in connectivity between the PHG and limbic network, with older adults demonstrating greater connectivity than younger adults. As a next step, we asked whether these mean connectivities were related to the magnitude of distinctiveness in the respective category-selective ROIs using Pearson correlations. The relationship between FG-visual network connectivity and distinctiveness in the FG was not significant (*r* = 0.14, *p* = 0.26).

Furthermore, the relationship between PHG-limbic network connectivity and distinctiveness in the PHG was not significant (*r* = -0.06, *p* = 0.64). No within-group relationships reached significance (*p*s > 0.16).

### 3.7 Interindividual variability in memory performance linked to regional distinctiveness

In order to understand whether age differences in distinctiveness and segregation were associated with memory, we performed Pearson correlations (see Figure 8). Memory performance was positively related to neural distinctiveness (*r* = 0.25, *p* = 0.04), suggesting that individuals demonstrating more distinctive neural representations also exhibited better recognition memory. The relationship between memory and distinctiveness was not significant within either age group (*p*s > 0.07) alone. Memory performance was further associated with network segregation (*r* = 0.26, *p* = 0.03), indicating that individuals with higher network segregation also tended to have better recognition memory. The relationship between memory and segregation was not significant within younger (*r* = 0.22, *p* = 0.20) or older (*r* = 0.25, *p* = 0.18) adults alone. No relationship was found between memory performance and connectivity between the FG and visual network (*r* = 0.01, *p* = 0.93).

**Figure 8.**
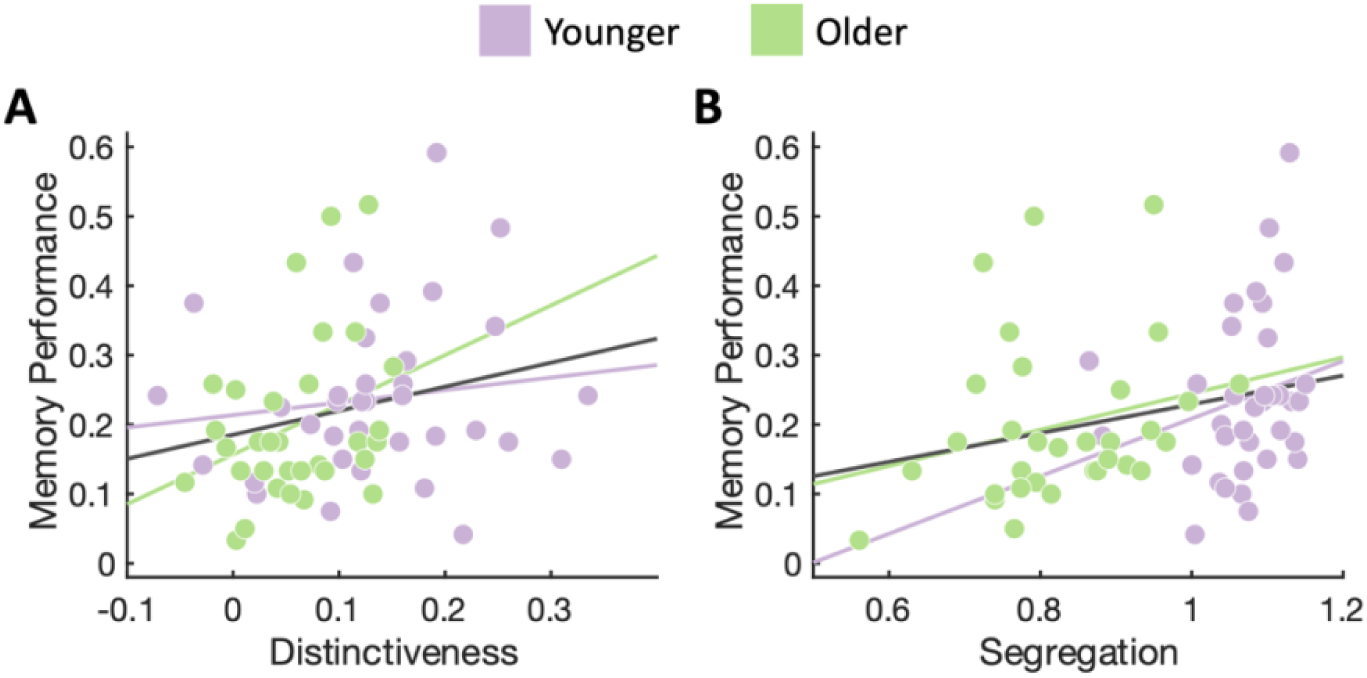
Scatter plots showing the relationship between memory performance and mean distinctiveness (A) and mean network segregation (B). Younger adults are shown in purple, older adults in green, and the whole sample in gray.

## 4. Discussion

Age-related neural dedifferentiation has garnered increasing attention over the last two decades as a potential underlying source of senescent cognitive decline (Li et al., 2001; for reviews, see Koen et al., 2020; Koen & Rugg, 2019; Sommer & Sander, 2022). Studies have reported evidence for neural dedifferentiation for individual stimuli (Koen, 2022; St-Laurent et al., 2014; Trelle et al., 2019), stimulus categories (Koen et al., 2019; Park et al., 2004; Pauley et al., 2023; Srokova et al., 2020), and global networks (Betzel et al., 2014; Chan et al., 2014; Geerligs et al., 2015; Varangis et al., 2019).

Thus far, few studies have investigated network dedifferentiation and looked for a link between network and regional dedifferentiation. Furthermore, whether aging targets the functional relationships of categorically-selective regions has yet to be explored. Here, in a sample of younger and older adults, we assessed age-related neural dedifferentiation across both the global network and categorical levels, their relation to one another, as well as the possibility that age differences in connectivity patterns to categorically-selective regions may be related to the observed regional dedifferentiation of categorical representations.

The present results showed that the neural distinctiveness of visual categories is reduced in older adults compared with younger adults, in line with the age-related neural dedifferentiation hypothesis (for reviews, see Koen and Rugg, 2019; Zhou et al., 2024). Specifically, we demonstrated that older adults exhibited lower representational specificity in the FG and PHG for faces and houses, respectively. Interestingly, age differences in neural distinctiveness were most consistent in the FG, with age differences observed during both encoding and recognition, compared to the PHG in which the age effect reached significance only during recognition. In contrast, prior work has suggested that age differences in distinctiveness tend to be more reliable for scene and house stimuli in the PHG compared to other stimulus categories, including faces and objects (Koen and Rugg, 2019; Srokova et al., 2020). Our findings indicate that age-related variability based on stimulus category may not be as straightforward as previously thought. For instance, it is difficult to disentangle whether age differences in neural distinctiveness in category-selective cortices are driven by stimulus properties specific to the stimulus category (e.g., stimulus complexity for scenes and houses; Garrett et al., 2020) or by region-specific effects (e.g., medial temporal cortices; Fjell et al., 2014). More studies are needed to understand how manifestations of age-related neural dedifferentiation vary across different stimulus categories.

Furthermore, we found evidence for an age-related disruption in intrinsic network organization, indicative of network dedifferentiation (see also, Koen et al., 2020). The observed age differences were driven by a combination of reduced within-network connectivity and increased between-network connectivity in older adults compared with younger adults in line with previous reports of dedifferentiated network structure in older age (Betzel et al., 2014; Cao et al., 2014; Chan et al., 2017, 2014; Geerligs et al., 2015; Iordan et al., 2017; King et al., 2018). Together, these findings implicate less segregated network organization as a robust feature of the aging brain.

Interestingly, age-related neural dedifferentiation of both category representations and functional networks was quite consistent across memory encoding and recognition, although age differences in network segregation were slightly stronger during recognition than during encoding. In a recent study using this same dataset, we uncovered a tight link between dedifferentiated category representations during encoding and retrieval, indicating that dedifferentiation in older adults during retrieval may be tied to either reinstatement of poorly encoded representations or poor perceptual processing during the recognition task (Pauley et al., 2023). The similarity in dedifferentiation across memory phases could further be attributed to the comparable attentional demands of the encoding and retrieval tasks in this study. Specifically, the target-detection task implemented during encoding used category-matched exemplars as targets, in which participants were required to focus on each stimulus in order to make a target judgment. This juxtaposes other, frequently-used target-detection tasks in which the fixation crosses are manipulated, allowing participants theoretically to make the target judgment without considering the task stimuli (e.g., Pauley et al., 2024; see Koen & Rugg, 2019 for further discussion). As such, here, the attentional resources needed during encoding were similar to those during retrieval, in which participants again attended each stimulus in order to make a memory judgment. In sum, this attentional overlap may have contributed to the consistencies in regional and network-level dedifferentiation across memory phases.

Additionally, regional dedifferentiation was associated with network-level dedifferentiation, but only when age was included in the model. Hence, this finding emphasizes that aging has a similar influence on both regional and whole-brain indices of neural function. Cassady and colleagues (2020) found a comparable relationship between distinctiveness of sensorimotor processing and segregation of the sensorimotor network. However, the findings from Cassady et al. (2020) were maintained regardless of age, leaving open the question as to whether regional distinctiveness and global network segregation might be related to one another. Together, these findings suggest that the age-related decline in distinctive neural processing appears to be a wide-ranging phenomenon that manifests similarly in both inter- and intraregional signaling, potentially as a result of an age-dependent physiological mechanism (Baltes and Lindenberger, 1997). For example, age-related dopaminergic dysregulation has been proposed to impair the fidelity of neural signaling and thus reducing distinctiveness in cortical representations (Li et al., 2000). In line with this proposal, dedifferentiation of neural representations has been linked to age differences in dopamine receptor concentrations (Abdulrahman et al., 2017). Additionally, blocking dopaminergic (D2) receptors reportedly mimicks age differences in functional network organization, albeit to a lesser extent (Achard and Bullmore, 2007). Thus, age differences in the dopaminergic system could potentially account for the related senescent declines in both regional distinctiveness and network segregation in the current study, though more work is needed to validate this theory.

Despite growing research focused on connectivity patterns to categorically-selective regions (Córdova et al., 2016; Furl, 2015; Keller et al., 2022; Norman-Haignere et al., 2012), the impact of age on these functional relationships has been so far overlooked. Here, we found the the FG was more strongly connected to the visual network compared to other networks and that the PHG was more strongly connected to the limbic network. Crucially, while we found an age-related increase in connectivity between the PHG and limbic network, we observed age-related declines in the coupling of the FG to the visual network, indicating a senescent disruption in connectivity between early and late visual cortices. Together, these findings suggest that task-relevant communication between visual cortices and the FG is impaired in older age.

Goh (2011) proposed that age-related reductions in distinctiveness of category-selective regions might be associated with impaired interregional communication. Although we identified both dedifferentiation of face representations in the FG and age differences in connectivity between the FG and visual network, we did not uncover evidence for a clear link between these findings. Thus, the question of whether regional dedifferentiation is tied to age differences in connectivity to category-selective cortices remains open. Future studies will be needed in order to determine whether or not a relationship exists between dedifferentiated category representations and the specific connectivity profiles of these category-selective cortices.

Finally, prior studies have underlined the importance of highly detailed representations to support memory processing across the adult lifespan (for reviews, see Koen et al., 2020; Koen and Rugg, 2019; Sommer and Sander, 2022; Zhou et al., 2024). These studies were additionally supported by findings suggesting that age differences in large-scale functional network organization coincide with an age-related reduction in mnemonic processing (see also, Chan et al., 2014; Persson et al., 2014). We found that interindividual variability in neural distinctiveness and network segregation were associated with memory performance across age groups, but not within age groups. This may be a limitation of the small sample size within each age group, but further studies should continue to investigate the relationship between age-related neural dedifferentiation and memory performance.

In summary, we find that older adults demonstrate declines in distinctiveness in categorical representations, connectivity to category-selective regions, and segregation of large-scale brain networks compared with younger adults. Due to the modest sample size, it is important to note that our findings should be interpreted as preliminary evidence, aiming to stimulate future research into the association between age-related neural dedifferentiation and age differences in functional network organization. Ultimately, understanding how functional neural representations differ between younger and older adults may be helpful to elucidate cognitive trajectories in older adulthood.

## Author statement

**Claire Pauley:** Formal analysis, Software, Writing - Original Draft. **Dagmar Zeithamova:** Methodology, Writing – Review & Editing. **Myriam C. Sander:** Conceptualization, Project administration, Supervision, Writing – Review & Editing.

## Acknowledgments

This work was conducted within the projects “Lifespan Age Differences in Memory Representations (LIME)” (PI: M.C.S.) and Lifespan Rhythms of Memory and Cognition (RHYME) (PI: Markus Werkle-Bergner) at the Max Planck Institute for Human Development. C.P. was a fellow of the International Max Planck Research School on the Life Course. D.Z. was supported by the National Institute of Neurological Disorders and Stroke (R01-NS-112366) as well as a Humboldt Research Fellowship for Experienced Researchers of the Alexander von Humboldt Foundation. M.C.S. was supported by the MINERVA program of the Max Planck Society. We thank all student assistants who helped with data collection, Gabriele Faust and members of the LIME and RHYME projects for helpful feedback, Julia Delius for editorial assistance, and all study participants for their time.

## References

Abdulrahman, H., Fletcher, P.C., Bullmore, E., Morcom, A.M., 2017. Dopamine and memory dedifferentiation in aging. NeuroImage 153, 211–220. 10.1016/j.neuroimage.2015.03.031

Achard, S., Bullmore, E., 2007. Efficiency and Cost of Economical Brain Functional Networks. PLOS Comput. Biol. 3, e17. 10.1371/journal.pcbi.0030017

Andrews-Hanna, J.R., Snyder, A.Z., Vincent, J.L., Lustig, C., Head, D., Raichle, M.E., Buckner, R.L., 2007. Disruption of large-scale brain systems in advanced aging. Neuron 56, 924–935. 10.1016/j.neuron.2007.10.038

Antonenko, D., Flöel, A., 2013. Healthy Aging by Staying Selectively Connected: A Mini-Review. Gerontology 60, 3–9. 10.1159/000354376

Baltes, P.B., Lindenberger, U., 1997. Emergence of a powerful connection between sensory and cognitive functions across the adult life span: A new window to the study of cognitive aging? Psychol. Aging 12, 12–21. 10.1037/0882-7974.12.1.12

Barense, M.D., Henson, R.N.A., Lee, A.C.H., Graham, K.S., 2010. Medial temporal lobe activity during complex discrimination of faces, objects, and scenes: Effects of viewpoint. Hippocampus 20, 389–401. 10.1002/hipo.20641

Betzel, R.F., Byrge, L., He, Y., Goñi, J., Zuo, X.-N., Sporns, O., 2014. Changes in structural and functional connectivity among resting-state networks across the human lifespan. NeuroImage 102 Pt 2, 345–357. 10.1016/j.neuroimage.2014.07.067

Cao, M., Wang, J.-H., Dai, Z.-J., Cao, X.-Y., Jiang, L.-L., Fan, F.-M., Song, X.-W., Xia, M.-R., Shu, N., Dong, Q., Milham, M.P., Castellanos, F.X., Zuo, X.-N., He, Y., 2014. Topological organization of the human brain functional connectome across the lifespan. Dev. Cogn. Neurosci. 7, 76–93. 10.1016/j.dcn.2013.11.004

Cassady, K., Gagnon, H., Freiburger, E., Lalwani, P., Simmonite, M., Park, D.C., Peltier, S.J., Taylor, S.F., Weissman, D.H., Seidler, R.D., Polk, T.A., 2020. Network segregation varies with neural distinctiveness in sensorimotor cortex. NeuroImage 212, 116663. 10.1016/j.neuroimage.2020.116663

Chan, M.Y., Alhazmi, F.H., Park, D.C., Savalia, N.K., Wig, G.S., 2017. Resting-State Network Topology Differentiates Task Signals across the Adult Life Span. J. Neurosci. 37, 2734–2745. 10.1523/JNEUROSCI.2406-16.2017

Chan, M.Y., Park, D.C., Savalia, N.K., Petersen, S.E., Wig, G.S., 2014. Decreased segregation of brain systems across the healthy adult lifespan. Proc. Natl. Acad. Sci. U. S. A. 111, E4997–E5006. 10.1073/pnas.1415122111

Chee, M.W.L., Goh, J.O.S., Venkatraman, V., Jiat, C.T., Gutchess, A., Sutton, B., Hebrank, A., Leshikar, E., Park, D., 2006. Age-related changes in object processing and contextual binding revealed using fMR adaptation. J. Cogn. Neurosci. 18, 495–507. 10.1162/jocn.2006.18.4.495

Chen, Q., Garcea, F.E., Almeida, J., Mahon, B.Z., 2017. Connectivity-based constraints on category-specificity in the ventral object processing pathway. Neuropsychologia 105, 184–196. 10.1016/j.neuropsychologia.2016.11.014

Chong, J.S.X., Ng, K.K., Tandi, J., Wang, C., Poh, J.-H., Lo, J.C., Chee, M.W.L., Zhou, J.H., 2019. Longitudinal Changes in the Cerebral Cortex Functional Organization of Healthy Elderly. J. Neurosci. 39, 5534–5550. 10.1523/JNEUROSCI.1451-18.2019

Córdova, N.I., Tompary, A., Turk-Browne, N.B., 2016. Attentional modulation of background connectivity between ventral visual cortex and the medial temporal lobe. Neurobiol. Learn. Mem. 134, 115–122. 10.1016/j.nlm.2016.06.011

Ebner, N.C., Riediger, M., Lindenberger, U., 2010. FACES-a database of facial expressions in young, middle-aged, and older women and men: Development and validation. Behav. Res. Methods 42, 351–362. 10.3758/BRM.42.1.351

Epstein, R., Kanwisher, N., 1998. A cortical representation the local visual environment. Nature 392, 598–601. 10.1038/33402

Esteban, O., Markiewicz, C.J., Blair, R.W., Moodie, C.A., Isik, A.I., Erramuzpe, A., Kent, J.D., Goncalves, M., DuPre, E., Snyder, M., Oya, H., Ghosh, S.S., Wright, J., Durnez, J., Poldrack, R.A., Gorgolewski, K.J., 2019. fMRIPrep: A robust preprocessing pipeline for functional MRI. Nat. Methods 16, 111–116. 10.1038/s41592-018-0235-4

Fair, D.A., Schlaggar, B.L., Cohen, A.L., Miezin, F.M., Dosenbach, N.U.F., Wenger, K.K., Fox, M.D., Snyder, A.Z., Raichle, M.E., Petersen, S.E., 2007. A method for using blocked and event-related fMRI data to study “resting state” functional connectivity. NeuroImage 35, 396–405. 10.1016/j.neuroimage.2006.11.051

Fjell, A.M., McEvoy, L., Holland, D., Dale, A.M., Walhovd, K.B., 2014. What is normal in normal aging? Effects of aging, amyloid and Alzheimer’s disease on the cerebral cortex and the hippocampus. Prog. Neurobiol. 117, 20–40. 10.1016/j.pneurobio.2014.02.004

Folstein, M.F., Folstein, S.E., McHugh, P.R., 1975. “Mini-mental state”. A practical method for grading the cognitive state of patients for the clinician. J. Psychiatr. Res. 12, 189–198. 10.1016/0022-3956(75)90026-6

Frank, L.E., Zeithamova, D., 2023. Evaluating methods for measuring background connectivity in slow event-related functional magnetic resonance imaging designs. Brain Behav. 13, e3015. 10.1002/brb3.3015

Furl, N., 2015. Structural and effective connectivity reveals potential network-based influences on category-sensitive visual areas. Front. Hum. Neurosci. 9, 253. 10.3389/fnhum.2015.00253

Garrett, D.D., Epp, S.M., Kleemeyer, M., Lindenberger, U., Polk, T.A., 2020. Higher performers upregulate brain signal variability in response to more feature-rich visual input. NeuroImage 217, Article 116836. 10.1016/j.neuroimage.2020.116836

Geerligs, L., Renken, R.J., Saliasi, E., Maurits, N.M., Lorist, M.M., 2015. A Brain-Wide Study of Age-Related Changes in Functional Connectivity. Cereb. Cortex 25, 1987–1999. 10.1093/cercor/bhu012

Goh, J.O.S., 2011. Functional dedifferentiation and altered connectivity in older adults: Neural accounts of cognitive aging. Aging Dis. 2, 30–48.

Gorgolewski, K.J., Auer, T., Calhoun, V.D., Craddock, R.C., Das, S., Duff, E.P., Flandin, G., Ghosh, S.S., Glatard, T., Halchenko, Y.O., Handwerker, D.A., Hanke, M., Keator, D., Li, X., Michael, Z., Maumet, C., Nichols, B.N., Nichols, T.E., Pellman, J., Poline, J.B., Rokem, A., Schaefer, G., Sochat, V., Triplett, W., Turner, J.A., Varoquaux, G., Poldrack, R.A., 2016. The brain imaging data structure, a format for organizing and describing outputs of neuroimaging experiments. Sci. Data 3, 1–9. 10.1038/sdata.2016.44

Gregorich, M., Strohmaier, S., Dunkler, D., Heinze, G., 2021. Regression with highly correlated predictors: Variable omission is not the solution. Int. J. Environ. Res. Public. Health 18. 10.3390/ijerph18084259

Haxby, J.V., Gobbini, M.I., Furey, M.L., Ishai, A., Schouten, J.L., Pietrini, P., 2001. Distributed and overlapping representations of faces and objects in ventral temporal cortex. Science 293, 2425–2430.

Iordan, A.D., Cooke, K.A., Moored, K.D., Katz, B., Buschkuehl, M., Jaeggi, S.M., Jonides, J., Peltier, S.J., Polk, T.A., Reuter-Lorenz, P.A., 2017. Aging and Network Properties: Stability Over Time and Links with Learning during Working Memory Training. Front. Aging Neurosci. 9, 419. 10.3389/fnagi.2017.00419

Kanwisher, N., McDermott, J., Chun, M.M., 1997. The fusiform face area: A module in human extrastriate cortex specialized for face perception. J. Neurosci. 17, 4302–4311. 10.1523/JNEUROSCI.17-11-04302.1997

Keller, A.S., Jagadeesh, A.V., Bugatus, L., Williams, L.M., Grill-Spector, K., 2022. Attention enhances category representations across the brain with strengthened residual correlations to ventral temporal cortex. NeuroImage 249, 118900. 10.1016/j.neuroimage.2022.118900

King, B.R., van Ruitenbeek, P., Leunissen, I., Cuypers, K., Heise, K.-F., Santos Monteiro, T., Hermans, L., Levin, O., Albouy, G., Mantini, D., Swinnen, S.P., 2018. Age-Related Declines in Motor Performance are Associated With Decreased Segregation of Large-Scale Resting State Brain Networks. Cereb. Cortex N. Y. N 1991 28, 4390–4402. 10.1093/cercor/bhx297

Kobelt, M., Sommer, V.R., Keresztes, A., Werkle-Bergner, M., Sander, M.C., 2021. Tracking age differences in neural distinctiveness across representational levels. J. Neurosci. 41, 3499–3511. 10.1523/jneurosci.2038-20.2021

Koen, J.D., 2022. Age-related neural dedifferentiation for individual stimuli: an across-participant pattern similarity analysis. Aging Neuropsychol. Cogn. 29, 552–576. 10.1080/13825585.2022.2040411

Koen, J.D., Hauck, N., Rugg, M.D., 2019. The relationship between age, neural differentiation, and memory performance. J. Neurosci. 39, 149–162. 10.1523/JNEUROSCI.1498-18.2018

Koen, J.D., Rugg, M.D., 2019. Neural dedifferentiation in the aging brain. Trends Cogn. Sci. 23, 547–559. 10.1016/j.tics.2019.04.012

Koen, J.D., Srokova, S., Rugg, M.D., 2020. Age-related neural dedifferentiation and cognition. Curr. Opin. Behav. Sci. 32, 7–14. 10.1016/j.cobeha.2020.01.006

Krishnan, A., Williams, L.J., McIntosh, A.R., Abdi, H., 2011. Partial Least Squares (PLS) methods for neuroimaging: A tutorial and review. NeuroImage 56, 455–475. 10.1016/j.neuroimage.2010.07.034

Li, S.-C., Lindenberger, U., Frensch, P.A., 2000. Unifying cognitive aging: From neuromodulation to representation to cognition. Neurocomputing 32–33, 879–890. 10.1016/S0925-2312(00)00256-3

Li, S.-C., Lindenberger, U., Sikström, S., 2001. Aging cognition: From neuromodulation to representation. Trends Cogn. Sci. 5, 479–486. 10.1016/S1364-6613(00)01769-1

Mak, L.E., Minuzzi, L., MacQueen, G., Hall, G., Kennedy, S.H., Milev, R., 2017. The Default Mode Network in Healthy Individuals: A Systematic Review and Meta-Analysis. Brain Connect. 7, 25–33. 10.1089/brain.2016.0438

Norman-Haignere, S.V., McCarthy, G., Chun, M.M., Turk-Browne, N.B., 2012. Category-Selective Background Connectivity in Ventral Visual Cortex. Cereb. Cortex 22, 391–402. 10.1093/cercor/bhr118

Park, D.C., Polk, T.A., Park, R., Minear, M., Savage, A., Smith, M.R., 2004. Aging reduces neural specialization in ventral visual cortex. Proc. Natl. Acad. Sci. U. S. A. 101, 13091–13095. 10.1073/pnas.0405148101

Park, J., Carp, J., Kennedy, K.M., Rodrigue, K.M., Bischof, G.N., Huang, C.M., Rieck, J.R., Polk, T.A., Park, D.C., 2012. Neural broadening or neural attenuation? Investigating age-related dedifferentiation in the face network in a large lifespan sample. J. Neurosci. 32, 2154–2158. 10.1523/JNEUROSCI.4494-11.2012

Pauley, C., Karlsson, A., Sander, M.C., 2024. Early visual cortices reveal interrelated item and category representations in aging. eNeuro 11, ENEURO.0337-23.2023. 10.1523/ENEURO.0337-23.2023

Pauley, C., Kobelt, M., Werkle-Bergner, M., Sander, M.C., 2023. Age differences in neural distinctiveness during memory encoding, retrieval, and reinstatement. Cereb. Cortex 33, 9489–9503. 10.1093/cercor/bhad219

Pauley, C., Sommer, V.R., Kobelt, M., Keresztes, A., Werkle-bergner, M., Sander, M.C., 2022. Age-related declines in neural selectivity manifest differentially during encoding and recognition. Neurobiol. Aging.

Payer, D., Marshuetz, C., Sutton, B., Hebrank, A., Welsh, R.C., Park, D.C., 2006. Decreased neural specialization in old adults on a working memory task. Neuroreport 17, 487–491. 10.1097/01.WNR.0000209005.40481.31

Persson, J., Pudas, S., Nilsson, L.-G., Nyberg, L., 2014. Longitudinal assessment of default-mode brain function in aging. Neurobiol. Aging 35, 2107–2117. 10.1016/j.neurobiolaging.2014.03.012

Power, J.D., Barnes, K.A., Snyder, A.Z., Schlaggar, B.L., Petersen, S.E., 2012. Spurious but systematic correlations in functional connectivity MRI networks arise from subject motion. NeuroImage 59, 2142–2154. 10.1016/J.NEUROIMAGE.2011.10.018

Pruitt, P.J., Tang, L., Hayes, J.M., Ofen, N., Damoiseaux, J.S., 2022. Lifespan differences in background functional connectivity of core cognitive large-scale brain networks. Neurosci. Res. 10.1016/j.neures.2022.09.005

Schaefer, A., Kong, R., Gordon, E.M., Laumann, T.O., Zuo, X.-N., Holmes, A.J., Eickhoff, S.B., Yeo, B.T.T., 2018. Local-Global Parcellation of the Human Cerebral Cortex from Intrinsic Functional Connectivity MRI. Cereb. Cortex 28, 3095–3114. 10.1093/cercor/bhx179

Silson, E.H., Steel, A., Kidder, A., Gilmore, A.W., Baker, C.I., 2019. Distinct subdivisions of human medial parietal cortex support recollection of people and places. eLife 8, e47391. 10.7554/eLife.47391

Snodgrass, J.G., Corwin, J., 1988. Pragmatics of measuring recognition memory: Applications to dementia and amnesia. J. Exp. Psychol. Gen. 117, 34–50. 10.1037/0096-3445.117.1.34

Sommer, V.R., Sander, M.C., 2022. Contributions of representational distinctiveness and stability to memory performance and age differences. Aging Neuropsychol. Cogn. 29, 443–462. 10.1080/13825585.2021.2019184

Song, J., Birn, R.M., Boly, M., Meier, T.B., Nair, V.A., Meyerand, M.E., Prabhakaran, V., 2014. Age-related reorganizational changes in modularity and functional connectivity of human brain networks. Brain Connect. 4, 662–676. 10.1089/brain.2014.0286

Srokova, S., Hill, P.F., Koen, J.D., King, D.R., Rugg, M.D., 2020. Neural differentiation is moderated by age in scene-selective, but not face-selective, cortical regions. eNeuro 7. 10.1523/ENEURO.0142-20.2020

St-Laurent, M., Abdi, H., Bondad, A., Buchsbaum, B.R., 2014. Memory reactivation in healthy aging: Evidence of stimulus-specific dedifferentiation. J. Neurosci. 34, 4175–4186. 10.1523/JNEUROSCI.3054-13.2014

Trelle, A.N., Henson, R.N., Simons, J.S., 2019. Neural evidence for age-related differences in representational quality and strategic retrieval processes. Neurobiol. Aging 84, 50–60. 10.1016/j.neurobiolaging.2019.07.012

Tzourio-Mazoyer, N., Landeau, B., Papathanassiou, D., Crivello, F., Etard, O., Delcroix, N., Mazoyer, B., Joliot, M., 2002. Automated Anatomical Labeling of activations in SPM using a macroscopic anatomical parcellation of the MNI MRI single-subject brain. NeuroImage 15, 273–289. 10.1006/nimg.2001.0978

Varangis, E., Habeck, C.G., Razlighi, Q.R., Stern, Y., 2019. The Effect of Aging on Resting State Connectivity of Predefined Networks in the Brain. Front. Aging Neurosci. 11. 10.3389/fnagi.2019.00234

Varangis, E., Habeck, C.G., Stern, Y., 2021. Task-based functional connectivity in aging: How task and connectivity methodology affect discovery of age effects. Brain Behav. 11. 10.1002/brb3.1954

Wang, X., Fang, Y., Cui, Z., Xu, Y., He, Y., Guo, Q., Bi, Y., 2016. Representing object categories by connections: Evidence from a mutivariate connectivity pattern classification approach. Hum. Brain Mapp. 37, 3685–3697. 10.1002/hbm.23268

Wolosin, S.M., Zeithamova, D., Preston, A.R., 2012. Reward modulation of hippocampal subfield activation during successful associative encoding and retrieval. J. Cogn. Neurosci. 24, 1532–1547. 10.1162/jocn_a_00237

Zhou, Q., Branton, G., Lessard, A., Polk, T.A., 2024. Dedifferentiation of neurocognitive function in aging, in: Reference Module in Neuroscience and Biobehavioral Psychology. Elsevier. 10.1016/B978-0-12-820480-1.00019-X

